# Deciphering the microbial architecture of pesticide and antibiotics biodegradation

**DOI:** 10.64898/2026.02.21.707176

**Authors:** Sylvia Thieffry, Julie Aubert, Jérémie Beguet, Marion Devers-Lamrani, Fabrice Martin-Laurent, Stéphane Pesce, Sana Romdhane, Nadine Rouard, Mathieu Siol, Aymé Spor

## Abstract

Understanding emerging functions at the scale of a bacterial community is a major challenge in microbial ecology and could lead up to promising tools for engineering microbial communities, for example in bioremediation. Here, through a top-down approach we obtained compositional variants of pesticide and antibiotics-degrading communities and further investigated communities features associated with their degradation abilities. We first tested whether diversity index or functional genes abundance could reliably be used as a proxy for this function, and obtained encouraging, albeit variable results. Further, through the use of statistical tools borrowed from the genomic selection literature, we were able to derive accurate prediction of the mineralisation potential of a bacterial community, based on its composition. However, the parallel between genotype-phenotype and community composition-mineralisation potential suffers a crucial caveat: bacterial abundances vary on a much wider scale than allele dosage at a given locus and are prone to change over time (particularly at the mineralisation scale). Here we observed that using presence/absence data instead of relative abundance can overcome these limitations and provide a clearer functional signal for mineralisation prediction through linear regression models. Random forest can also intrinsically deal with microbial data without transformation and select for significant predictors. We suggest drawing inspiration from the tools and concepts used in genotype-phenotype mapping to elucidate microbial functions at the community level while keeping in mind the significant differences between these two fields. This parallel is here exemplified by the concept of microbial architecture of degrading functions, akin to the genetic architecture of phenotypic traits.

## Introduction

Microbial communities are essential for ecosystem functioning and, as such, are drawing increasing attention for biotechnological purposes in relation to ecosystem health (Kostic et al., 2024). Even though their use is centuries old, a fine understanding of biological processes they perform made possible by recent technological advances opened new doors for harnessing their wide potential. The targeted manipulation of microbial communities toward a feature of interest, known as microbiome engineering or microbial community design, is still an emerging field of science (Albright et al., 2021; Matuszyńska et al., 2024; Rillig et al., 2015; Silverstein et al., 2023). Among the numerous functions performed by microbes, their ability to degrade and convert xenobiotic compounds into nutrients is of prime interest and can be translated into bioremediation approaches to improve ecosystem health. However, to date, the successful establishment of a degradation function following a single microbial strain inoculation has been hampered by many biotic and abiotic barriers (Lopes et al., 2022; Thompson et al., 2005). This calls for a change of scale, shifting from the use of isolated strains to more complex and resilient microbial consortia or engineered microbial communities where the relationship between microbial communities compositions and structures and their degradation capacity is further elucidated (Pileggi et al., 2020).

One can address the functional understanding of a process with a bottom-up approach, based on mechanistic and *a priori* knowledge. In line with genetic engineering, the focus is on metabolic processes at the sub-individual scale of enzymes or genes, often with an aim of process modelling (San Román et al., 2025). Here, information on microbial degradation of the two xenobiotics of interest, namely isoproturon (IPU), a phenyl-urea herbicide, and sulfamethazine (SMZ), a sulfonamide antibiotic, is easily available from the literature.

The only IPU microbial mineralisation pathway identified to date involves *pdmAB* genes, coding for an N-demethylase, in turn responsible for the initial step of IPU degradation in various *Sphingomonads* (Gu et al., 2013; Hussain et al., 2015; Sørensen et al., 2003). In *Sphingomonads*, these genes are located on a plasmid, suggesting that horizontal gene transfer plays a role in IPU-degrading genes mobility. In addition, as many catabolic genes of xenobiotics, *pdmAB* genes are flanked by insertion sequences suggesting that mobile genetic elements can play important roles in the evolution of IPU-mineralisation ability (Yan et al., 2016). As such, the abundance of the *pdmA* gene could be a more precise indicator of the degradation capacity than the abundance of known degrading bacteria, as recently observed in a microcosm experiment (Storck et al., 2020).

On the other hand, sulphonamide-catabolising genes *sadABC* or close homologues have been identified more recently (Ricken et al., 2017) and found in various strains (Cao et al., 2019; Kim et al., 2019; Reis et al., 2018; Ricken et al., 2017; Wu et al., 2023). A recent metagenomic study reports *sadABC* genes in members of *Micrococcaceae* and *Microbacteriaceae* (Chen et al., 2023). Little is known about their genetic mobility, but as they are surrounded by insertion sequences and found in several copies in several strains, it suggests potential homologous recombinations of *sad* genes in various genomic environments (Kim et al., 2019; Ricken et al., 2017). Rogue et al. showed a good correlation between SMZ biodegradation potential and abundance of the *sadA* gene in communities, as previously observed for IPU (Rogue et al., 2025).

Conversely, a top-down approach is based at the community scale and does not require mechanistic insights. As such, a global index of community diversity could be informative about functional performance. Indeed, complementary effects (use of diverse resources) and/or selection effects (more chances to contain species with large functional effects) are expected to generate a positive link between community diversity and function (Loreau, 2000).

At a finer scale, species presence and abundance can be used to link microbial composition and structure to a specific community function. This emergent property is somewhat analogous to the emergence of complex phenotypic traits based on eukaryotes genetic information (Morris et al., 2020; Sanchez et al., 2023). While genome-wide association studies attempt to decipher the genetic make-up of a trait by identifying statistical associations between genetic markers along the genome and the expression of a particular phenotype, genomic prediction further uses this rationale to derive predictions of phenotypes from the genetic composition. Genetic architecture, referring to “the landscape of genetic contribution to a given phenotype” (Timpson et al., 2017) *i*.*e*. the number of genes influencing the phenotype and their effect sizes, as well as their interactions, can be very different depending on the phenotype (e.g. human height is very polygenic, while the soapy taste of cilantro is controlled by a single-nucleotide polymorphism (Eriksson et al., 2012)). In the same vein, in microbial communities, highly integrative “phenotypes” (functional traits) such as biomass production or respiration are more likely to be influenced by a majority of the species in the community. By contrast, specific phenotypes such as xenobiotic degradation are more likely driven by the presence and abundance of a few bacterial groups. Even more so than in the previous genetics analogy, interactions between microbes are likely to lead to departures from the simple additive model where the phenotype results from the simple addition of the parts’ phenotypic effects (Diaz-Colunga et al., 2024).

Leveraging the genotype/phenotype and community composition/function analogy, statistical methods from genomic selection can be applied to microbial communities with the aim to decipher the architecture of xenobiotic degradation and predict degradation potential from community composition. Recent studies using penalised regression and machine-learning methodologies have already shown promising results for various community traits (Bien et al., 2021; Hinton and Mucha, 2021; Statnikov et al., 2013; Thompson et al., 2019).

Here, using a top-down manipulation approach, through the combination of dilutions and biocidal treatments of two xenobiotic-degrading source communities, we explored the community/function landscape (San Román et al., 2025) and set out to link community composition to degradation capacity.

## Materials and methods

### Environmental sampling

Enhanced bacterial degradation of various xenobiotics has been reported in different environments regularly exposed to these compounds (James et al., 2010; Topp et al., 2013). Therefore, we collected two “environmental matrices” with previous exposure to xenobiotic compounds: (i) an arable soil with a historical use of IPU and (ii) a river sediment recurrently exposed to urban contamination, including pharmaceuticals such as SMZ. The arable soil was collected from Epoisses INRAE experimental farm (Bretenière, France, 47°30 ‘22.18”N, 4°10 ‘26.46”E) with the following properties: 51.9% silt, 41% clay and 6.2% sand, pH 7.2, organic carbon content 15.5 g/kg dry soil and nitrogen 1.4 g/kg. The soil was pre-incubated in mesocosms at IPU agronomical dose (2 mg/kg dry soil). The sediment was collected from the downstream of Tillet river (Aix-les-Bains, France, 45°41’44.10”N, 5°53’44.76”E, see (Rogue et al., 2025) for more details) with the following properties: 3.2% silt, 9.4% clay and 95.9% sand, pH 8.6, organic carbon content 5.7 g/kg dry soil and nitrogen 0.2 g/kg. It was pre-incubated for six months in an aquarium with monthly SMZ treatment (nominal concentration of 100 mg/kg dry weight) to promote and maintain degradation capacities. The soil and sediment were sieved to 4 mm, and a sufficient amount of each matrix was sterilised by *γ*-irradiation for further use (2 times 35 kGy in Ionisos, Dagneux, France).

### Experimental design

The experimental procedure to produce community variants and analyse their mineralisation capacity and composition is presented in figure 1. Soil and sediment communities, which respectively exhibited IPU- and SMZ-degrading capacities, were extracted by blending 33 g equivalent dry mass of either soil or sediment with 60 mL sterile distilled water or 60 mL of supernatant, respectively. The D1 dilution was obtained by diluting soil and sediment suspensions 10 times. To lower diversity levels by removing the rare community members, D1 was diluted 100 times to give D2 and D3 was obtained following a final 10-fold dilution of D2.

We coupled each dilution with 8 biocidal treatments to further modify community composition: three pH conditions by addition of acids and bases followed by washes (malic acid 1 M for pH2, malic acid (10^−2^M) for pH4, ammonia 20% 1 M for pH11), oxidative treatments by addition of H_2_O_2_ 0.98 M (13 µl for mild and 27 µL for strong.), heat stress at 70°C, UV exposure for 2h and xenobiotic treatments (IPU for SMZ-degrading communities and reciprocally). 210 µL from resulting suspensions were inoculated into microcosms containing 1 g dry soil or sediment, in 24 wells plate as follows: IPU-degrading variants plus controls (n=4), were inoculated into sterile sediment, except the “matrix control” samples (n=4) that were inoculated back into sterile soil (while SMZ-degrading variants were inoculated into sterile soil, except the matrix controls that were inoculated back into sterile sediment). Soil moisture was adjusted to 60% of WHC (Water-Holding Capacity) while 1 mL sterile water was added to sediment samples to mimic waterlogged environmental conditions.

Samples were treated with IPU at a nominal concentration of 2 mg/kg dry matrix (agronomical dose) or SMZ at a nominal concentration of 1 mg/kg dry matrix (approximately the maximum concentration found in rivers (Ducrocq et al., 2024)), with ^14^C-xenobiotic addition in half of the samples. Xenobiotic mineralisation was measured by radiorespirometry on samples with radioactivity while the others were kept incubating in the dark at 21°C. When mineralisation reached a plateau (35 days for SMZ-degrading communities and 66 days for IPU-degrading communities), community composition at the OTU level was assessed on non-radioactive replicates.

### Assessing mineralisation capacities by radiorespirometry

Xenobiotic mineralisation was measured by micro-radiorespirometry following ISO 14239:2017. Briefly, a mixture of ^12^C-xenobiotic (Sigma) and ^14^C-phenyl-labelled marked xenobiotic (with specific activity of 0.67 GBq/mmol for IPU and 0.015 MBq/mmol for SMZ) to reach 1300 Bq radioactivity was added to 1 g of soil/sediment dry weight equivalent per sample. Then samples were incubated in 24-well plates (BioLite, Thermo Fisher Scientific) in the dark at room temperature (20°C) until the mineralisation dynamics reached a plateau. Plates were covered with blotting paper (Chromatography Paper, Whatman R, England) soaked in barium hydroxide solution (from crystallised octahydrate, Sigma, in sterile water to reach maximum solubility) which traps emitted CO_2_. For each time point, blotting paper were replaced and ^14^CO_2_ quantity determined by the use of a phosphor screen (Molecular Dynamics), optical scanner (Storm 860 Molecular Imager, Molecular Dynamics) and image analysis software (ImageQuant^™^ TL 10.2, Cytiva). Mineralisation dynamics were expressed as the cumulative percentage of ^14^CO_2_ evolved from the initially added ^14^C-xenobiotic over time.

Degradation capacities were analysed based on three distinct features estimated from the mineralisation kinetics: (i) the total mineralisation at the end of mineralisation dynamics (cumulative percentage of ^14^C-xenobiotic mineralised as ^14^CO_2_). (ii) the initial mineralisation rate during the first days of mineralisation dynamics (1/3 of the total of days) expressed in percentage of mineralised ^14^C-xenobiotic per day. (iii) the membership to a group of mineralisation based on the characteristics of the entire mineralisation dynamics (time before start mineralisation, mineralisation rate, total, time to reach a plateau). The clustering of the samples based on their mineralisation kinetics into groups was performed using functional data analysis (R package fda 6.1.8). This consists of a PCA (Principal Component Analysis) on functional data followed by an AHC (Ascending Hierarchical Classification) on euclidean distances between smoothing factors. Number of groups was based on Ward.D2 criterion of relative inertia loss.

### Sequencing microbial community

250 mg equivalent dry mass of each matrix were retrieved from all 192 samples to extract DNA using DNeasy PowerSoil-htp 96 well DNA isolation kit (Qiagen, Hilden, Germany). Generation of amplicons for Illumina MiSeq sequencing was done using a two-steps PCR protocol according to Berry et al. (Berry et al., 2011). Amplification in duplicate of the V3-V4 hyper-variable region of the bacterial 16s rRNA was done with fusion primers U341F (5’-CCTACGGGRSGCAGCAG-3’) and 805R (5’-GACTACCAGGGTATCTAAT-3’), and, to allow the subsequent addition of multiplexing index sequences, overhang adapters were used (forward: TCGTCGGCAGCGTCAGATGTGTATAAGAGACAG, reverse: GTCTCGTGGGCTCGGAGATGTGTATAAGAGACAG). All samples were then cleaned with SequalPrep Normalization plate kit 96-well (Invitrogen, Carlsbad, CA, USA), pooled and sent for sequencing on MiSeq (Illumina, 2 × 250 bp) using the MiSeq reagent kit v2 (500 cycles).

The sequence data were analysed using an in-house developed Jupyter Notebook (Kluyver et al., 2016) piping together different bioinformatics tools. Briefly, R1 and R2 sequences were assembled using PEAR (Zhang et al., 2013) with default settings. Further quality checks were conducted using the QIIME pipeline (Caporaso et al., 2009) and short sequences were removed (< 400 bp). Reference based and *de novo* chimera detection, as well as clustering in OTUs were performed using VSEARCH (Rognes et al., 2016) and the adequate reference databases (SILVA’ representative set of sequences). The identity threshold was set at 94% based on replicate sequencing runs of a bacterial mock community containing 40 bacterial species.

### Microbial diversity and community composition

All statistical analyses were carried out using R version 4.4.2. Five communities (2 for IPU, 3 for SMZ) were pre-filtered and set aside due to low number of reads and/or total dominance of a specific OTU, probably caused by technical difficulties. *α*-diversity was measured through Simpson’s Reciprocal Index, using the microbiome package (Leo Lahti, 2019). For further analysis, OTUs were filtered based on prevalence, keeping those at least present (frequency > 0.0001%) in 6% of samples, lowering counts to 883 OTUs in IPU communities and 748 OTUs in SMZ communities. Principal Coordinates Analyses (PCoAs) were performed on Aitchison distance (Gloor et al., 2017), calculated after zero imputation through a Bayesian-multiplicative replacement procedure using the cmultRepl function from the zCompositions package (Palarea-Albaladejo and Martin-Fernandez, 2015).

### Quantification of xenobiotic degrading genes

Real-time PCR reactions were carried out according to the ISO 17601 standard in a ViiA7 thermocycler (Life Technologies, United States) in a 15 µL reaction volume. The total bacterial community was quantified using 16S rRNA gene primers described by Muyzer et al. (Muyzer et al., 1993). SMZ- and IPU-degrading communities were quantified using *sadA* primers described by Billet et al. (Billet et al., 2021) and *pdmA* primers described by Storck et al. (Storck et al., 2020). For each real-time PCR reaction, the mixture contained 7.5 µL of Takyon MasterMix (Eurogentec, France), 1 µM of each primer, 250 ng of T4 gene 32 protein (QBiogene, France) and 2 ng of DNA. Thermal cycling conditions were set as follows: 95°C for 3 min, 35 cycles of 95°C for 30 s, annealing temperature (depending on the considered gene primers) for 30 s and 72°C for 30 s. Each real-time PCR assay was performed twice independently in order to check repeatability. Standard curves were realised with serial dilutions of linearised plasmids containing the appropriate cloned target genes (ranging from 0.5*e*2 to 0.5*e*7 number of gene copies per µl). The mean values of efficiency are 98.3% for 16S rDNA, 92.4% for *pdmA* gene and 67.7% for *sadA* gene. Melting curves were visually inspected to verify the amplification specificity for each gene. The absence of qPCR inhibitors was also tested by “house-developed” method, using plasmid DNA (blank pGEM-T Easy Vector, Promega, France) as a control.

Diversity and qPCR data were subjected to statistical analysis of variance using a Kruskal-Wallis test followed by Dunn’s test for multiple comparisons (Bonferroni adjusted p-value *≤* 0.05) using the rstatix package (version 0.7.2).

### Predicting degradation function from microbial community diversity

We aimed at predicting degradation features (total mineralisation, initial mineralisation rate and mineralisation group membership) based on OTU abundance. Unlike genomic data, OUT counts are compositional data (Gloor et al., 2017) and that has to be taken in consideration before model training. With that in mind, we tested two different data transformations, each of them supporting an underlying hypothesis on OTUs’ impact on community degradative function:

i. Total Sum Scaling (TSS): OTUs counts are divided by total sample reads. The underlying hypothesis supports the idea that abundance data is important for prediction.
ii. Presence/absence (Pres/Abs): OTU counts are replaced with 1 if >0 and let at 0 otherwise, hypothesizing that presence is informative enough for prediction.

All the following analyses are based on three statistical methods used in genomic prediction: ElasticNet (EN), LASSO, and Random Forest (RF), which were implemented in the glmnet version 4.1-8 and ranger version 0.16.0 R packages, respectively. EN and LASSO are penalised regression techniques suitable for situations with many predictors (OTUs) and few observations (“large p, small n”). They help to reduce the variance of estimates, for LASSO by selecting variables, and EN by both selecting and shrinking them. RF, a machine-learning method based on decision trees, does not assume additive effects among OTUs, making it effective for modelling complex, non-additive and non-linear relationships.

Predictions were computed using the tidymodels framework (R package version 1.2.0, (Kuhn and Wickham, 2020). The first step is to split the data into a training and a testing set, respectively 3/4 and 1/4 of the data, stratified according to the predicted variable (here the mineralisation measurement). A 10-fold cross validation scheme with five repetitions was used on the training set to tune hyper-parameters: the penalty parameter for penalised regression, plus a selection-shrinkage balance parameter for EN, and node size and variables number per split for RF with number of trees set to 500. Models were then trained to minimize the RMSE (root mean square error) or, in the case of predicting group membership, to maximize AUC ROC (Area Under Curve of the Receiver Operating Characteristic). We then assessed the quality of prediction on the testing set, either by looking at these metrics or by computing Pearson’s correlation between predicted and actual data. The whole process was repeated eight times, redrawing the sets each time. Variable importance was retrieved using vip package 0.4.1.

### Co-occurrence networks

For each of the two xenobiotics (IPU and SMZ), a bacterial co-occurrence network was inferred using a zero-inflated sparse Poisson lognormal model (R package PLNmodels v1.2.0) (Batardière et al., 2024; Chiquet et al., 2019), with a fixed zero-inflation parameter. The best model was selected based on a Stability Approach to Regularisation Selection (Liu et al., 2010) with a stability parameter of 0.99 (high stability). Networks manipulation and visualisation were made using R package igraph v2.0.3 (Csardi and Nepusz, 2006). Louvain method from the same package (based on modularity optimization) was used to extract community structure of networks, discriminating bacteria into “Louvain groups” (instead of so-called Louvain communities, to avoid confusion).

## Results

### Impact of dilutions and treatments on bacterial community diversity, structure and functions

The two degrading communities were subjected to dilution and biocidal treatments for creating community variants with expected contrasting IPU- and SMZ-degradation capacities. We transferred these community variants and control communities (CT, no biocidal treatments) into a new sterile matrix, *i*.*e*., in sediment for IPU-degrading communities and soil for SMZ-degrading communities, with the exception of CTM controls (back into their initial sterilised matrix). Bacterial community composition of all communities was then assessed for each replicate when the mineralisation dynamics of the two compounds reached a plateau.

Comparing CT and CTM samples, we evaluated the effect of environmental filtering on the transfer of degradation function between soil and sediment (Fig. 2 A&B). SMZ-degrading communities derived from sediment showed similar mineralisation potential when inoculated in soil (CT) compared to sediment (CTM) (e.g. 22.5% *±*1.6 and 21.6% *±*1.5 for D1) but were sensitive to dilution, with 22.5% *±*1.6 in D1, 14.7% *±*1.5 in D2 and 11.8% *±*1.6 in D3 for CT. IPU-degrading communities were not affected by the environmental change from soil (CTM) to sediment (CT) when the entire community was inoculated (even rising from 32.9%*±*0.3 in CTM to 59.1%*±*2.9 in CT for D1). In contrast, while dilution significantly impacted IPU mineralisation potential in sediments, down to 0.6% *±*0.2 for CT-D3, it tended to have the opposite effect when diluted control communities were inoculated back into soil, with up to 47.7% *±*1.8 for CTM-D3.

The effect of dilution was also clear when considering biocidal treatments, with, overall, a higher mineralisation achieved in D1 for both IPU and SMZ communities across treatments. Some specific treatments, namely pH2, pH11 and the heat stress (HS), resulted in the total loss of mineralisation potential even for D1 dilutions with mineralisation below 5% for IPU (for pH2, pH11 and the HS) and for SMZ (HS only). However, the combined effect of dilution and treatments seemed to be more progressive in the case of SMZ-degrading communities with mineralisation potential decreasing gradually from D1 to D3, while in the case of IPU-degrading communities, the response of the mineralisation potential was more abrupt. Overall, these results indicate that changing environments (i.e. from soil to sediment or vice versa) is not a strong barrier for transferring SMZ and IPU degradation capacities provided that the transferred communities are not too impoverished.

Dilutions coupled with biocidal treatments also resulted in significant differences in *α*-and *β*-diversity of bacterial communities (Fig. 2 C, D, E & F). Biocidal treatments had a strong negative impact on *α*-diversity of IPU-degrading community, compared with the mild effect on SMZ-degrading community (Fig. 2 C & D). However, this does not reverse the trend towards greater diversity in IPU communities, especially in controls.

As expected, diluting communities lowered their *α*-diversity (Fig. 2 C & D). In IPU-derived communities, this was mainly due to the loss of less abundant OTUs (SI Fig. 1A (Thieffry et al., 2026)), while in SMZ-derived communities, it was primarily driven by changes in OTU relative abundances, as the number of observed OTUs remained within the same range across samples (SI Fig. 1B (Thieffry et al., 2026)). Moreover, dilutions is a driver in modifying community composition as seen in PCoA (Fig. 2 E & F), with communities clustering depending on dilution across treatments.

**Figure 1.**
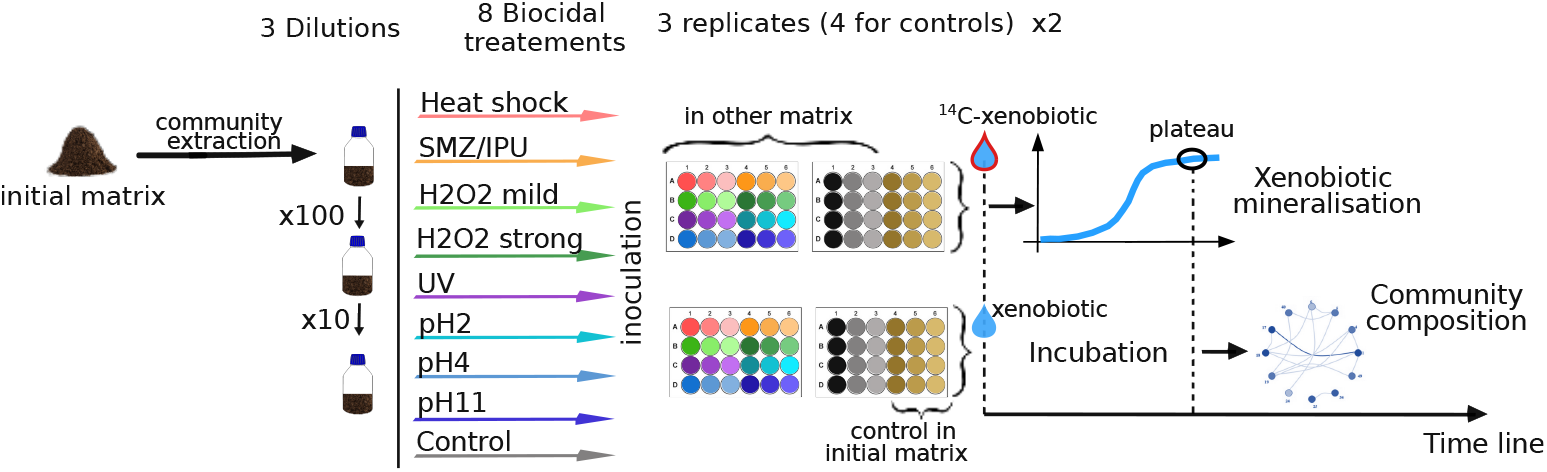
Experimental procedure to engineer community variants and evaluate their mineralisaton capacities and community compositions. The degrading community underwent several dilutions coupled with biocidal treatments, then resulting community variants were inoculated into a new sterile matrix with addition of ^14^C-xenobiotic. Mineralisation was measured by radiorespirometry while community composition (at the OTU level) was assessed after the mineralisation plateau was reached.

**Figure 2.**
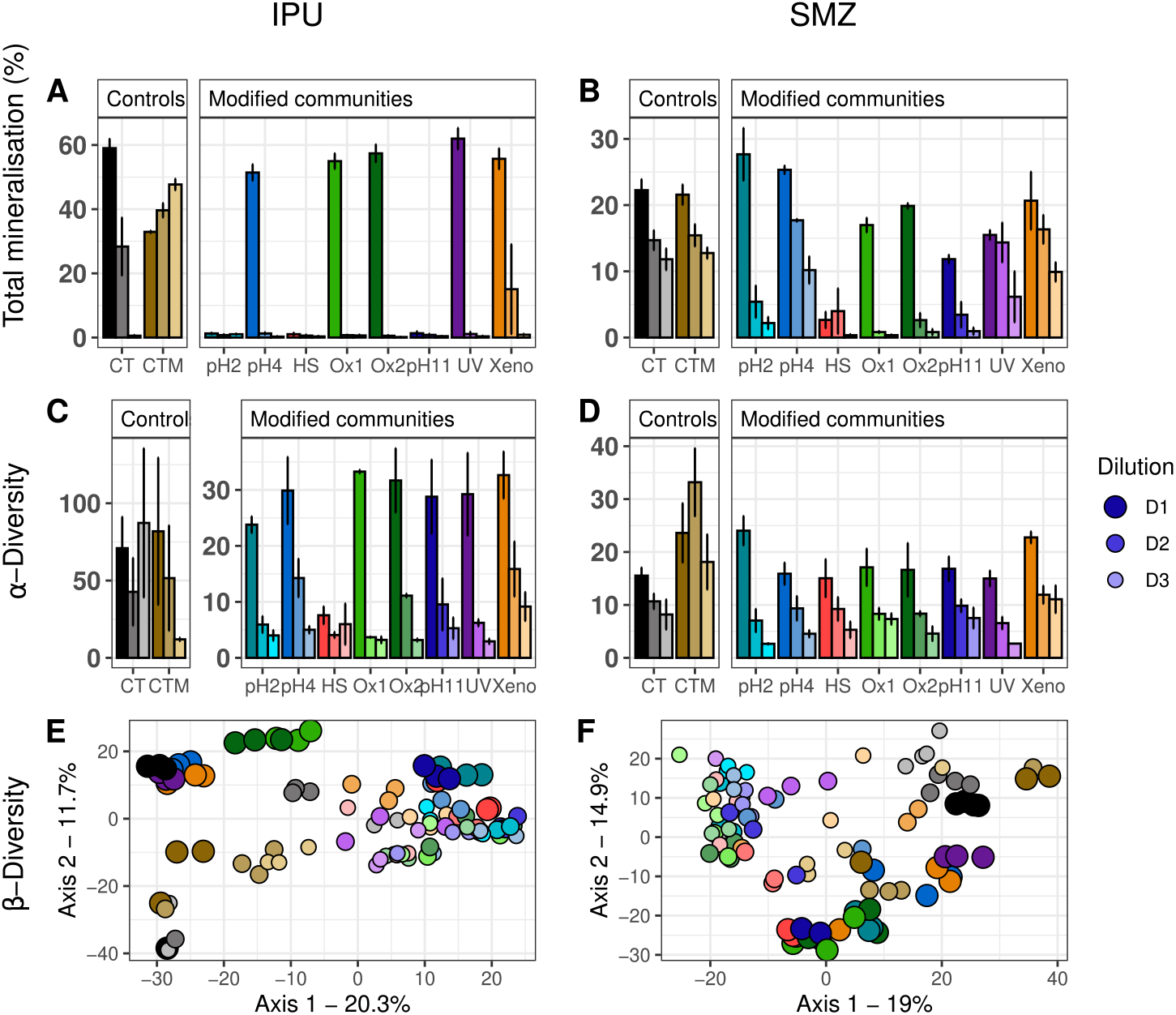
Impact of dilutions and treatments on IPU (left) and SMZ (right) mineralisation potential and associated *α*-and *β*-diversity of bacterial communities. Three dilutions (D1, D2 and D3) and eight treatments (pH2, 4 and 11; HS, Ox 1 and 2; UV; xeno) were tested on two initial matrices (soil and sediment). Barplots correspond to one condition of biocidal treatments (x-axis, in different colors) and one dilutions (as illustrated by the blue gradient in the legend). Controls: communities inoculated in initial matrix (CTM) and different matrix (CT). The rest of community variants are all in different matrix. (A & B) Mineralisation potential expressed as the percentage of ^14^CO_2_ evolved from the ^14^C-xenobiotic initially added. Error bars show standard error of the mean with n = 3, with exception of controls n = 4. (C & D) Simpson’s Reciprocal Index of communities. (E & F) PCoA on Aitchison distance with same colour code as in A, B, C, D.

### Linking biodegradation capacities to diversity levels and degrading genes quantification

We further classified all the samples based on their mineralisation dynamics into three groups using functional data analysis tool (Fig. 3 A & C). For IPU-degrading compositional variants, 62 were affiliated to group 1 (G1), comprising non-IPU-mineralising samples (mean of 1.15% mineralisation), 23 were affiliated to group 2 (G2) with high IPU-mineralising samples (mean of 55.40%) and 11 were affiliated to group 3 and were found in between (mean 37.97%) (Fig. 3 A).

**Figure 3.**
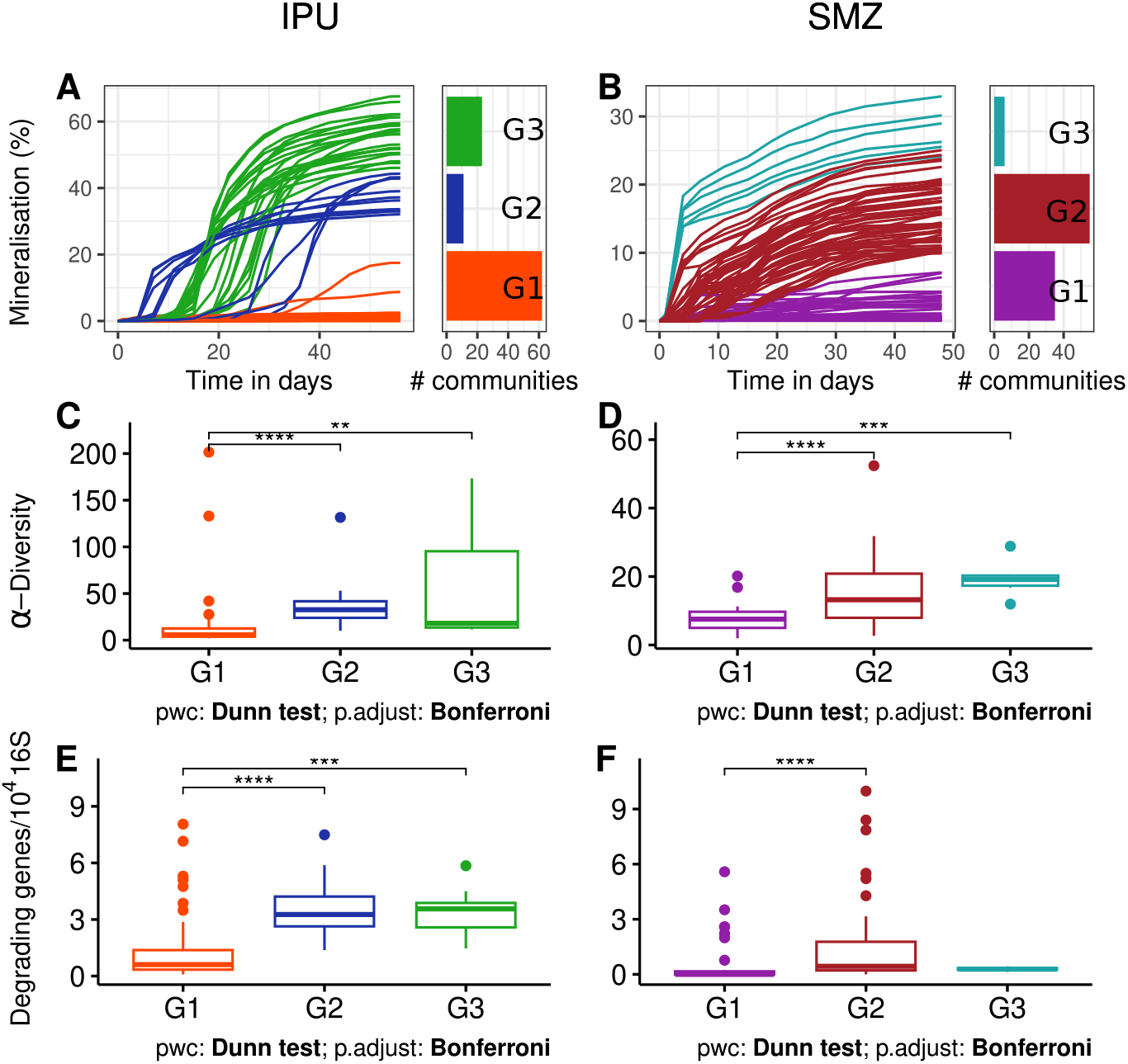
Characteristics of the three mineralisation groups for IPU (left) and SMZ (right) degrading communities. (A & B) Mineralisation kinetics clustered in three groups distinct by colour (G1, G2 and G3) followed by community counts by groups. (C & D) Diversity shown as Simpson’s Reciprocal Index by degrading groups. (E & F) Ratio of known degrading genes on 16S by degrading groups. For IPU (E), relative abundance of *pdmA* genes and for SMZ (F), relative abundance of *sadA* genes.

Regarding SMZ, lower mineralisation levels were reached and a disparity between groups was observed. Six samples mineralised fast, reaching a mean of 28.01% total mineralisation (G3), 35 did not mineralise well with a mean of 1.95 % (G1) and the rest, 55 samples, were found in between, averaging 15.75% of ^14^C-SMZ mineralisation (G2). We further investigated the bacterial community diversity of these groups and quantified the abundance of xenobiotic-specific degrading genes, *pdmA* and *sadA* for IPU and SMZ, respectively.

For IPU-degrading communities, diversity in G1 (14.49*±*3.80) was significantly lower than G2 (36.02*±*5.23) and G3 (57.98*±*20.10; Dunn test with Bonferroni adjusted p<0.05). For SMZ, G1 communities (7.68*±*0.68) were also less diverse than G2 (15.13*±*1.25) and G3 (5.54*±*2.26; Dunn test with adjusted p<0.05). These results reinforced our hypothesis regarding the importance of diversity for degrading ability.

As anticipated, the abundance of the *pdmA* gene (Fig; 3 E), responsible for the first and key step of the bacterial transformation of IPU, is linked to degradation capacities, discriminating degrading groups as well as does the diversity level (1.29 *×* 10^−4^ *±* 2.1 *×* 10^−5^ *pdmA* per 16S rDNA as mean of G1 compared to 3.58 *×* 10^−4^ *±* 3.1 *×* 10^−5^ and 3.39 *×* 10^−4^ *±* 3.6 *×* 10^−5^ respectively means of G2 and G3).

Regarding SMZ, *sadA* abundance (Fig. 3 F) was significantly lower in G1 non-mineralising communities (5.7*×*10^−5^ *±*2.1*×*10^−5^ *sadA* per 16 rDNA) compared to G2 (1.37*×*10^−4^ *±*2.2*×*10^−5^), able to better mineralise SMZ. However, surprisingly, the best mineralising group, G3, as well as several samples of G2 (SI Fig. 2 (Thieffry et al., 2026)) displayed a relatively low abundance of *sadA* gene (G3 : 2.9 *×* 10^−4^ *±* 4.3 *×* 10^−6^), suggesting the existence of another SMZ-degrading pathway.

### Predicting degrading capacities based on community composition

The next step consisted in linking xenobiotic degradation potential represented by three parameters (total mineralisation, initial rate of mineralisation on the first days and degradation group) to the composition of bacterial community at the OTU level. This was done comparing the performance of three prediction and classification statistical methods, namely Random Forest (RF), Elastic net (EN) regression and LASSO regression. Two data transformations were also tested for their intrinsic differences and high performances (Karwowska et al., 2025) : Total Sum Scaling (TSS) or presence/absence (PresAbs) of the predictive variables (see SI Fig. 4 (Thieffry et al., 2026)).

**Figure 4.**
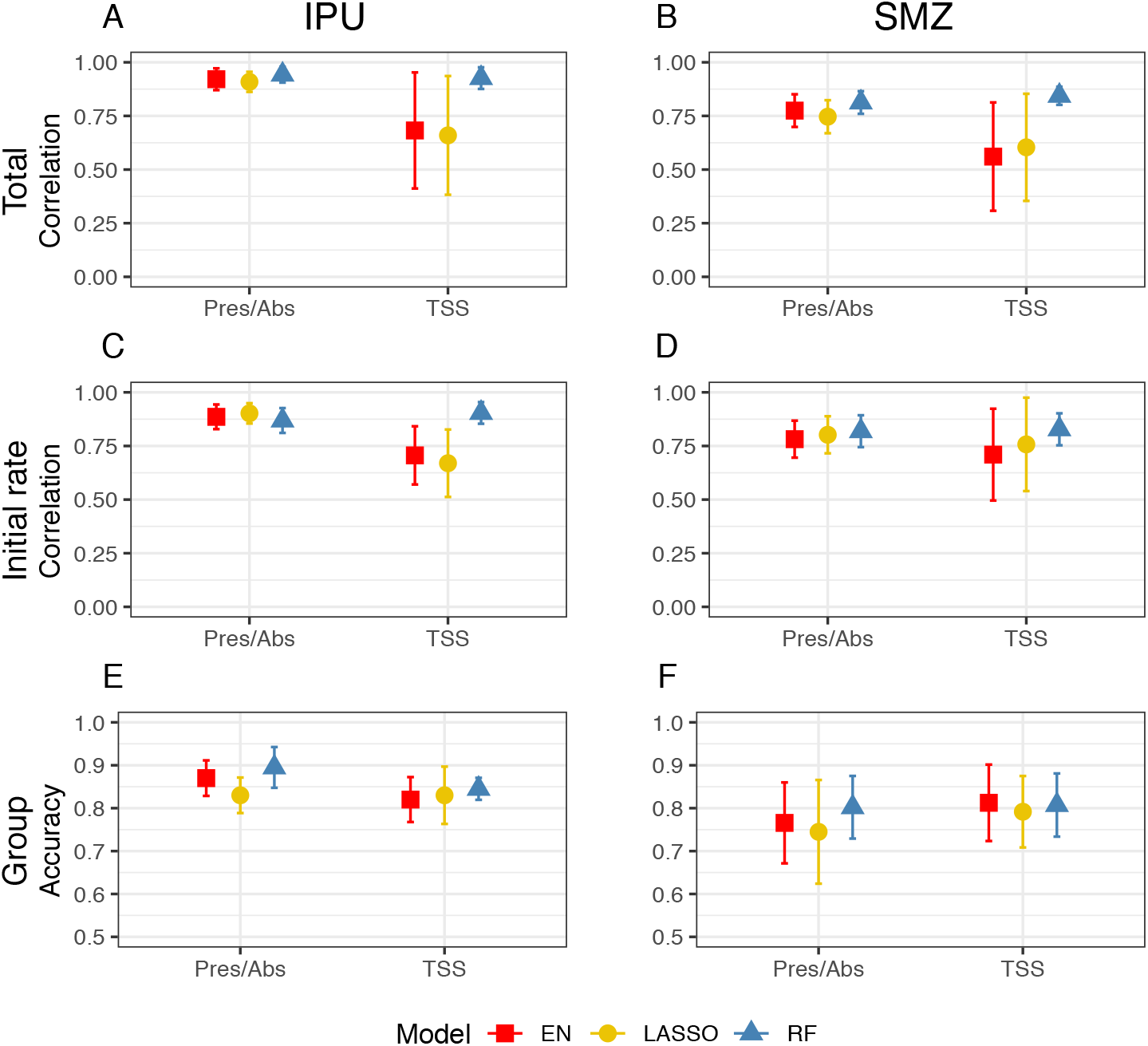
Correlation between measured and predicted degradation parameters and classification accuracy for mineralising groups. Pearson’s correlation between actual mineralisation measurements (Total, Initial rate) and values predicted by elastic net regression (EN), LASSO, and random forest (RF) on relative abundance OTUs data (TSS) and presence Absence of same OTUs (Pres/Abs). Accuracy of mineralising group (Group) classification by same statistical methods. All predictions were assessed by 10-fold CV with shuffling between eight runs, and only results on test set are shown. Error bars stand for the standard error of the mean.

Correlation between true and predicted values of Total and Initial rate of mineralisation are depicted in Fig. 4 (A to D). First, we observe overall better results in the case of IPU (half of correlations above 0.9) than for SMZ degradation (no correlation above 0.9). Moreover, the PresAbs transformation yielded better and less variable correlation results for regression models compared to TSS (e.g. for Total IPU mineralisation correlations are 0.92*±*0.02 and 0.91*±*0.02 in PresAbs and 0.68*±*0.10 and 0.66*±*0.10 in TSS). A similar pattern was found for SMZ, while the correlation on RF prediction is not sensitive to the OTUs transformation (e.g. 0.94*±*0.01 in PresAbs and 0.93*±*0.02 for Total IPU mineralisation). No clear difference can be established between the two regression method performances, even if LASSO tended to be slightly inferior than Elastic Net, particularly when classifying in groups with different degradation profiles (SI Fig. 7 (Thieffry et al., 2026)). Besides, neither the classification method nor data transformation seem to impact the accuracy of prediction when trying to assign group membership to the communities. Concerning IPU groups, accuracy stands around 0.85, with particular difficulties in predicting membership to G3 (e.g. see SI Fig. 5 & SI Fig. 6 (Thieffry et al., 2026)). This could be explained by the small size of the group, as well as the similarity with G2-type curves (Fig. 3). The same situation of unbalanced group composition for SMZ community in addition to the lack of between-group disparity results in lower classification accuracy when looking at SMZ samples.

### Predictive OTUs and community network

Next, we examined the OTUs with higher predictive value (*i*.*e*., with larger effect sizes), by selecting the top 25 OTUs for each mineralisation measure/data and transformation/model combinations (SI Fig. 8 & 9 (Thieffry et al., 2026)). Predictive values of OTUs derived from occurrence in decision trees (RF) are always positive and can be considered analogous to linear coefficients, positive or negative, from linear regression. To have a global look at their distribution depending on model and data transformation, we illustrated the scaled absolute value of the best predictive values for the two xenobiotics (Fig.5) and found an interesting pattern, similar for both compounds.

Surprisingly, OTUs identified using TSS and regression models were amongst the least abundant ones, with ranks lower than 200 (with one exception) and a large majority in the last third (Fig. 5), while it was not the case for OTUs identified by PresAbs and Random Forest. This finding was consistent across mineralisation measures (Total, Initial rate, Group) and xenobiotics, even though opposite to our expectations. This pattern is in line with the lower prediction capacities of regression models based on OTUs relative abundance (TSS), even though less clearly in the case of group prediction (Fig. 4).

**Figure 5.**
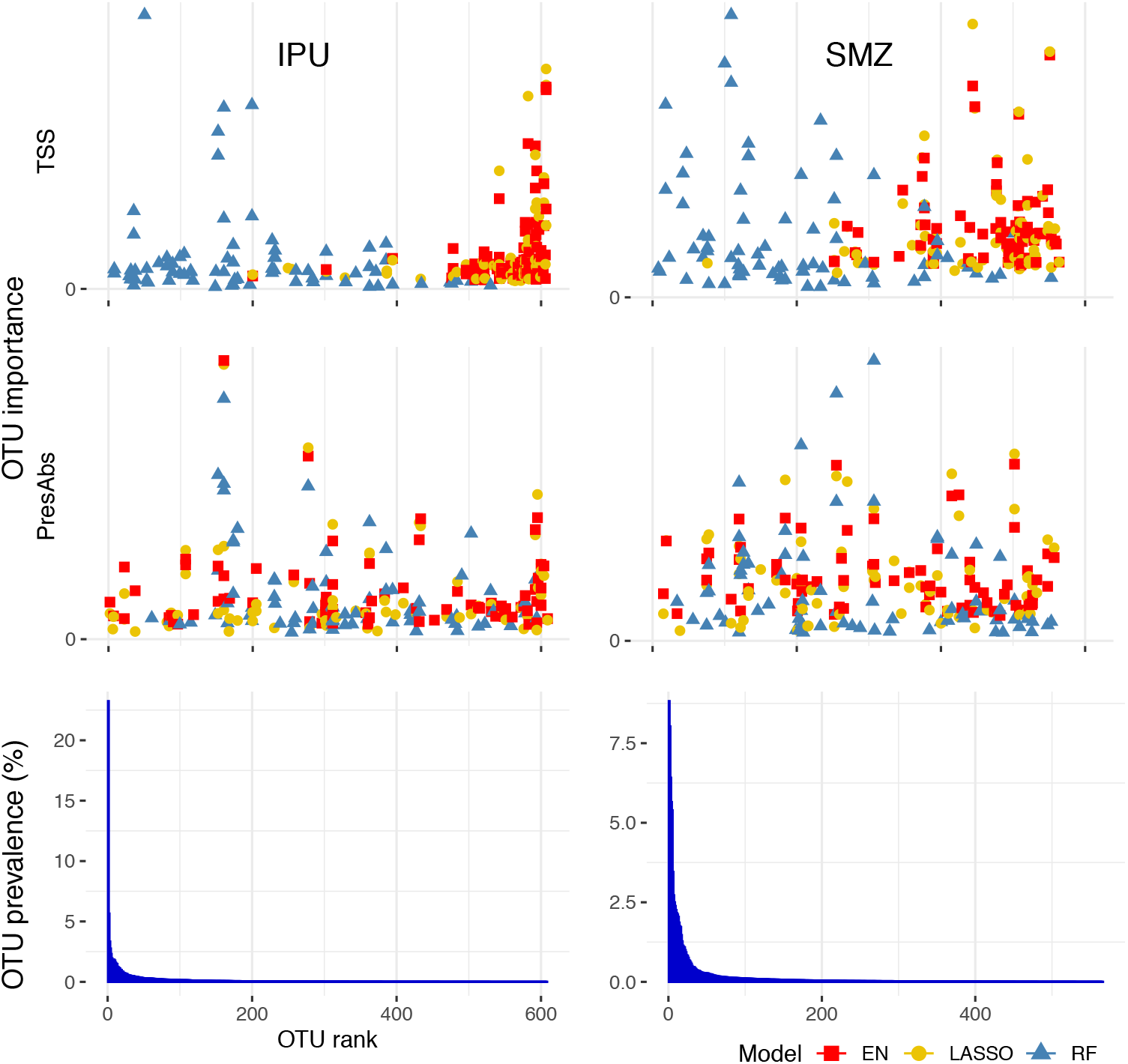
OTU predictive value. OTUs are displayed by decreasing abundance rank (same axis for the three plots). Relative abundance is expressed in percent across all samples. Scaled predictive values of top 25 OTUs for each mineralisation measures and associated prediction/classification methods are shown (different colours and shapes for EN, LASSO models and RF). OTU predictive values in PresAbs were retrieved from model based on presence/absence of OTUs while TSS from model based on their relative abundance.

Moreover, predictive OTUs identified by LASSO and EN were largely overlapping (transformation and mineralisation measures wise) while OTUs important for Random Forest (RF) predictions were mostly different (SI Fig. 10 & Fig. 11 (Thieffry et al., 2026)). We therefore pooled predictive OTUs from LASSO and EN for further analyses. Interestingly, important OTUs identified by RF were more frequently shared across transformation (TSS and PresAbs, 1/3 shared) compared to LASSO or EN (less than 1/10), underlying the fact that RF is less affected by the choice of the data transformation.

To further investigate predictive OTUs, we constructed co-occurrence network for each xenobiotic using a zero-inflated sparse multivariate Poisson log-normal model. From these networks, we selected OTUs with predictive power in regression models after TSS transformation (Fig. 6 A&D) and PresAbs transformation (Fig. 6 B&E) and their immediate neighbours (cooccurring OTUs). Resulting networks are far smaller when based on TSS OTUs, with, for SMZ, 14 OTUs from regression and 54 links of co-occurrence compared to PresAbs where 29 OTUs from regression are inserted in the network, forming 133 links. A similar pattern is found in IPU-degrading community, with 16 main OTUs forming 80 links in TSS while 30 OTUs are forming 159 links in PresAbs.

**Figure 6.**
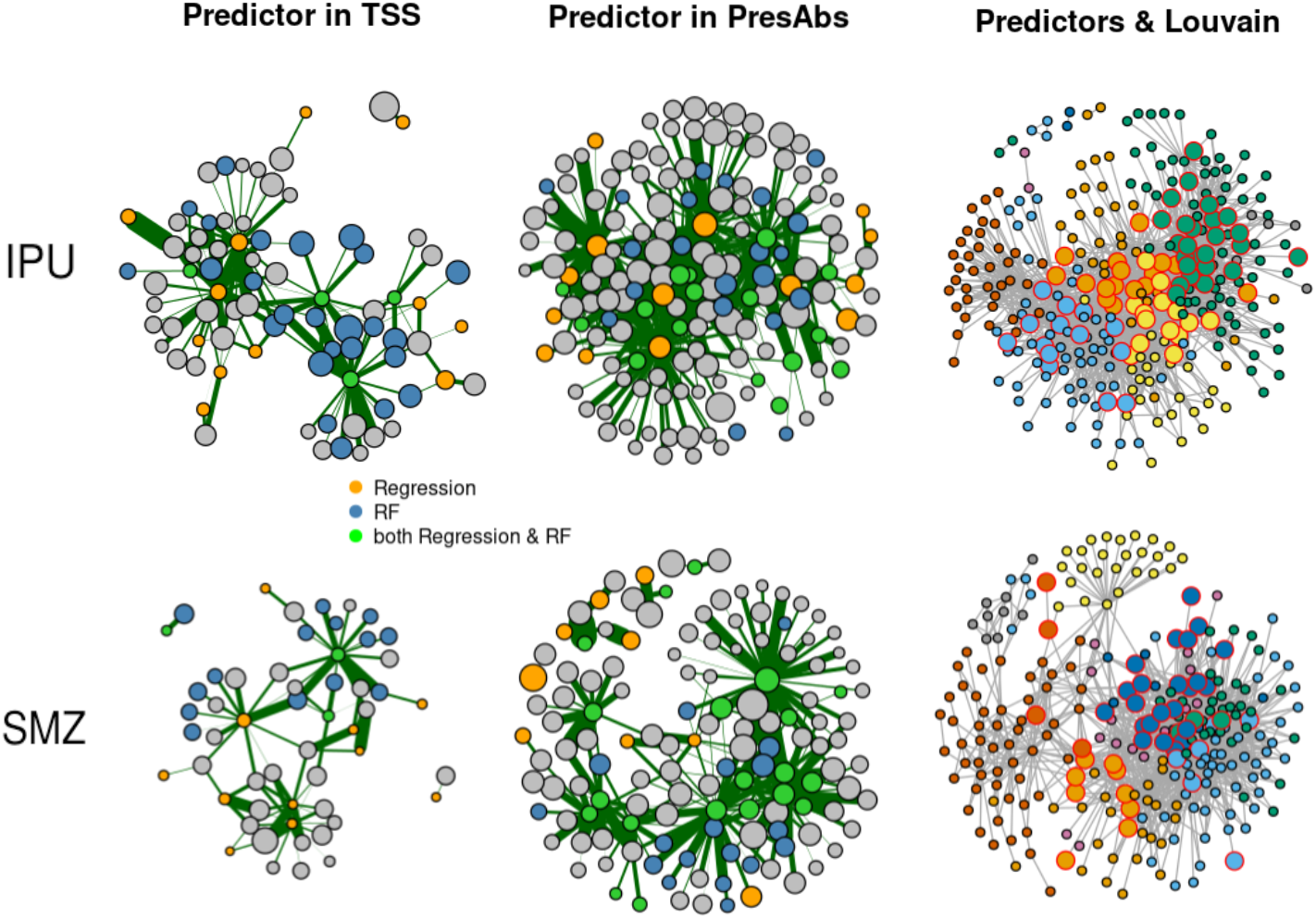
Co-occurrence networks & Predictive OTUs. Bacterial network inferred from OTU abundances across treatments and separately for each of the two xenobiotics. A, B, D & E show OTU predictors from regression models (in orange) and their direct cooccurring OTUs (in grey), highlighting those that are predictors for RF prediction in blue and the ones that we found in regression and RF in green. Edge thickness is proportional to partial correlation, positive in green and negative (extremely rare) in red. Size of the nodes is proportional to the logarithm of the total OTU abundance across samples. Entire co-occurrence network are represented in E & F, with nodes coloured by Louvain groups. Larger nodes encircled in red are OTU predictors common between TSS and PresAbs and their direct co-occurring OTUs predictive either in TSS or PresAbs.

We then highlighted OTUs predictive in RF (blue nodes) and common between the two methods -RF and regression- (in green), showing that they are largely intertwined. More precisely, for networks in TSS, low-abundance OTUs predictive in regression are co-occurring with a great number of abundant OTUs important for RF predictions with few overlaps of the two (IPU : 25 only for RF, 4 predictive for both RF and regression; SMZ : 14 RF and 3 overlaps). In contrast, there is a large overlap for PresAbs predictors between RF and regression (IPU : 21 RF and 16 overlaps, 18 RF and 20 overlap for SMZ).

Predictive OTUs appeared well linked to each other for the majority, with the exception of 8 predictive OTUs in the SMZ network. To confirm this proximity, we used the Louvain method to detect groups by modularity optimisation on the overall co-occurrence network, and pointed out OTUs with predictive value in both TSS and PresAbs with no model preference. The selected OTUs have attested predictive power but, when aiming at interpreting this as degrading capacities, one should also consider closely linked OTUs, as co-occurrent OTUs have by definition similar distribution and prediction models choose indiscriminately one or another. For this reason, we also highlighted co-occurrent OTUs with predictive value in one of the mineralization measures/data transformation/model combinations. They are all depicted in 6 C&F, coloured by Louvain group membership : 9 groups for SMZ-degrading communities, with 5 prevalent (more than 20 OTUs) and 11 for IPU of which 5 are prevalent. Predictive OTUs are exclusively part of prevalent groups for both xenobiotics, implying that they are found among the most connected part of community (SI Fig. 12 (Thieffry et al., 2026)).

## Discussion

In this study, our aim was to provide a proof of concept exploring the link between community and function, using pesticide (IPU) and antibiotic (SMZ) degradation as model functions. Using a top-down manipulation approach combining dilutions and biocidal treatments applied to initial communities from two different environments (namely soil and sediment), we studied IPU and SMZ degradation functions by assessing the importance of various community characteristics such as diversity, abundance of xenobiotic-degrading genes, environmental change, and finally community composition at the OTU level.

Environmental filtering has been shown to be a major factor for shaping bacterial community diversity and structure in soil (Calderón et al., 2016; Yan et al., 2019). As such, it is a possible first hurdle to overcome when aiming at introducing a bacterial community in a new environment. Here we tested the introduction of various xenobiotic-degrading communities in a contrasting recipient environment from soil to sediment or the other way around, and found no strong abiotic hindrance regarding the targeted function (Albright et al., 2021; Graham et al., 2016). Perhaps counter-intuitive, this result could be explained by the niche offered by the xenobiotic treatment, facilitating the establishment of the degrading communities. Only the introduction into sediment of IPU-degrading communities that had reached a low level of diversity due to dilution led to a loss of function. However, since dilution also causes a decrease in bacterial density, it cannot be ruled out that this is also involved in the loss of function. In any case, these results support the hypothesis that the diversity and density of the inoculum drive the outcome of an invasion process (Huet et al., 2025; King et al., 2023; Silverstein et al., 2023; Vila et al., 2019).

Our top-down community manipulation approach successfully resulted in the production of a range of variants presenting different diversity levels, allowing us to test our working hypothesis of a positive link between microbial diversity and community function. This association has been empirically assessed by previous work on widespread community functions such as respiration or carbon use (Bell et al., 2005; Langenheder et al., 2010; Yu et al., 2019). These authors noted the importance of complementarity mechanisms in successfully performing functions, even though, at very high diversity, the ubiquitous functional redundancy effect could cause this relation to saturate (Langenheder et al., 2010). In our case, the specificity of IPU and SMZ degradation pathways argues rather for a selection effect: there are more chances for diverse communities to include degrading bacteria. However, uniform level of species richness in SMZ-degrading communities (SI Fig. 1 (Thieffry et al., 2026)) tempers this hypothesis and underlines the importance of community evenness for functional performance. That way, complementarity mechanisms could be of importance when looking at specific functions via facilitative interactions. Overall, community diversity gives relevant but qualitative information on xenobiotic degradation, differentiating only very low degradation levels from higher ones.

The quantification of *pdmA* genes in IPU-derived communities allowed us to discriminate between degrading and non-degrading communities with nearly the same level of precision as when using a diversity index such as Simpson’s reciprocal (3). Yet what we have here is essentially a qualitative signal with no room for quantitative predictive power, incurring a relatively high false positive rate, with many communities showing high abundance of *pdmA* genes, yet no mineralisation potential (SI Fig. 2E (Thieffry et al., 2026)). Concerning SMZ degradation, the gene-function link is even looser in our results, with no clear association between the mineralisation performance and the amount of *SadA* amplicons (SI Fig. 2F (Thieffry et al., 2026)). A possible explanation for this would be that we miss a part of the picture and there are other genes, not targeted in our study and not yet known, enabling SMZ degradation. Timing of gene analysis is also of prime importance, as shown by several temporal studies on xenobiotic mineralisation : abundance of degrading genes increases only during the exponential degradation phase before falling back to or under detection level (Martin-Laurent et al., 2003; Storck et al., 2020). It is likely that our qPCR analysis was too late to efficiently track degrading genes, with levels of degrading genes too close to the detection threshold specifically in the SMZ case, leading to low reliability in the data. Our results highlight pitfalls when deducing functional information based on gene abundance, supporting Rocca et al. conclusions. Their work challenged the gene abundance-function relationship by looking at various microbial processes (N and C cycles, xenobiotic degradation) and finding no convincing link to genomic data (Rocca et al., 2014).

Next, the parallel between genetic information/phenotype and community composition/degradation capacities led us to apply statistical methods from genomic selection on microbial community composition data (OTU abundance) to predict degradation capacities. The prediction results highly depend on data transformation and the chosen method. We found no significant differences between LASSO and Elastic Net predictions, with predictive power shared by only a small number of OTUs in both cases (SI Fig. 8 & 9 (Thieffry et al., 2026)), suggesting that there is no need for information from many predictors, corresponding to a relatively oligogenic architecture in genetic term (oligos = few, little, describing a trait influenced by a few genes). Moreover, significant predictors from linear regression models are showing contrasted abundance profiles depending on data transformation, with almost exclusively low abundant OTUs chosen from TSS data. The best results are obtained with Random Forest (RF) with no influence of data transformation as well as regression methods on presence/absence data.

From there, we can assume that linear regression models do not easily handle microbiota data due to high sparsity and great variability (typical OTU distribution consists of many zeros and some very high counts). The cause can lie in the technical effect of the imprecision of OTU quantification when based on 16S rRNA gene (Brooks et al., 2015; Louca et al., 2018). On the biological side, it could be explained by a threshold effect of OTU predictors, with no proportional effect above some specific threshold count. Also, temporal dynamics can have a prevailing part in this: on the one hand, degrading bacteria abundance can substantially change over time and on the other hand community analysis can be performed at different times regarding the degradation process itself, which could result in a drastically weakened link between OTUs relative abundance and total degradation or degradation rate. For classification into mineralisation groups, however, the information used is the shape of the entire mineralisation kinetics, and as such could be less influenced by the timing of the measurement, resulting in a better agreement between predictions based on PresAbs data vs TSS data. We presume that we depart from these identified issues when simplifying data from relative abundance to PresAbs with linear regression models or when using random forest. Indeed, in these cases we obtained higher correlations between predicted and measured degradation values and less prediction variability. These overall outperformance of models based on PresAbs (or equivalent results for RF) are in line with recent findings and recommendations from classification of human phenotypes (healthy/diseased) based on microbiomes (Giliberti et al., 2022; Karwowska et al., 2025).

OTUs selected as predictors by linear regression based on presence absence and RF are largely overlapping or co-occurring across communities, displaying common sources of information. This also underlies that accurate predictions can rely on co-occurring OTUs, even if they are not identified as mineralising OTUs. Furthermore, co-occurrence of OTUs may reflect biologically relevant links such as biotic interactions but could also indicate similar niche preferences without direct interaction (Blanchet et al., 2020; Goberna and Verdú, 2022). Because of this limited biological interpretability, prediction results can only provide limited insights into degradation pathways, identification of functional guilds or partition of the degradation process. Still, we show that degradation capacities can be reliably predicted from a set of biomarkers, which consistently belong to a small number of modular groups in bacterial communities.

## Conclusion

The comprehension of ecological functions performed by microorganisms at the community level is a promising tool for harnessing their wide potential. Through the example of xenobiotic biodegradation we tested whether diversity indices or functional gene abundance could reliably be used as a proxy for this function, and obtained encouraging, albeit variable results. Further, through the use of statistical tools borrowed from the genomic selection literature, we were able to derive accurate prediction of the mineralisation potential of a bacterial community, based on its composition. However, the parallel between genotype/phenotype and community composition/mineralisation potential suffers a crucial caveat: bacterial abundances vary on a much wider scale than allele dosage at a given locus and are prone to change over time (particularly at the mineralisation scale). Here we observed that using presence/absence data instead of relative abundance can overcome these limitations and provide a clearer functional signal for mineralisation prediction through linear regression models. Random Forests can also intrinsically deal with microbial data without transformation and select significant predictors. These predictive bacteria are well inserted in the microbial community and further research is needed to determine if they only are biomarkers or if they play a key role in the mineralisation process. Finally, our work advocates for the application of ideas from genotype-phenotype mapping on microbial community data with the aim to unravel microbial functions at the community scale.

## Acknowledgements

We thank members of the EMFEED team for their help and support.

## Data, script, code, and supplementary information availability

Data, script and codes as well as supplementary information are available online (https://doi.org/10.5281/zenodo.20396547; Thieffry et al., 2026). Raw sequences were deposited at the NCBI under the BioProject PRJNA1418749.

