## Supplementary Figures for "Deciphering the microbial architecture of pesticide and antibiotics biodegradation" (from another version of this preprint: version 1, 2026-02-23)

February 2, 2026

#### 1 Alpha diversity - Observed species of degrading communities

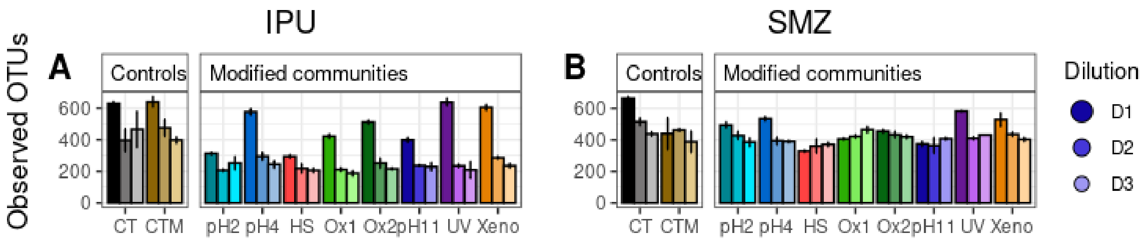


Figure 1: **Observed species diversity for IPU (left) and SMZ (right) degrading communities.** Barplots correspond to one condition of biocidal treatments (x-axis, in different colours) and one dilutions (colour gradient). Error bars are standard error of the mean with n = 3, with exception of controls n = 4. Controls: communities inoculated in initial matrix (CTM) and different matrix (CT), like all modified communities

#### 2 Total mineralisation on diversity and degrading genes

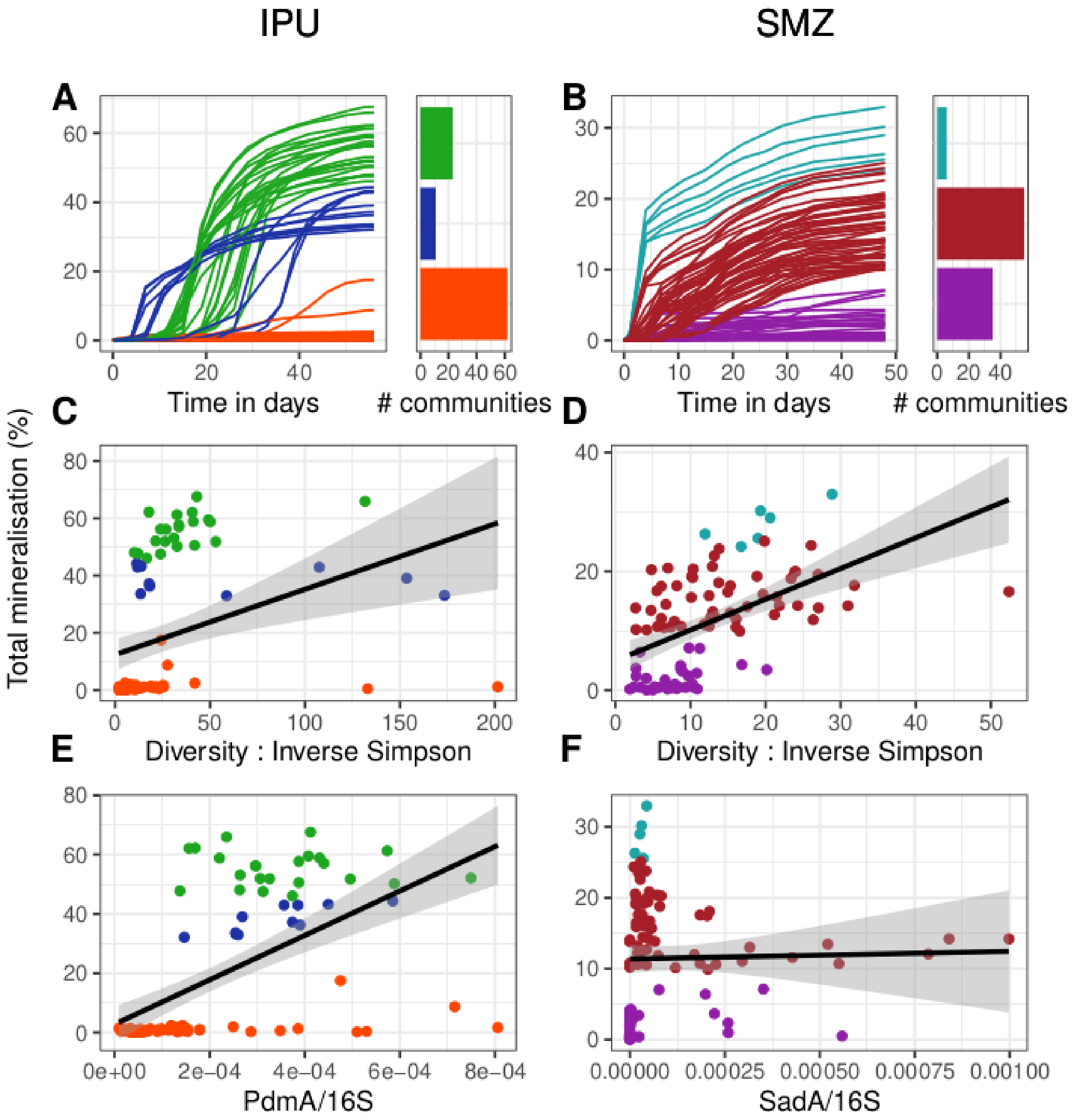


Figure 2: **Mineralisation on IPU- (left) and SMZ (right)- degrading communities features.** (A & B) Mineralisation dynamics clustered in colour distinctive groups followed by community count by groups. (C & D) Total mineralisation on diversity shown as Simpson's Reciprocal Index. (E & F) Total mineralisation on ratio of known degrading genes on 16S. For IPU (E), *pdmA* genes and for SMZ (F) *sadA* genes. C to F, linear regression is shown with standard error.

#### 3 Error (RMSE & AUROC) of prediction models

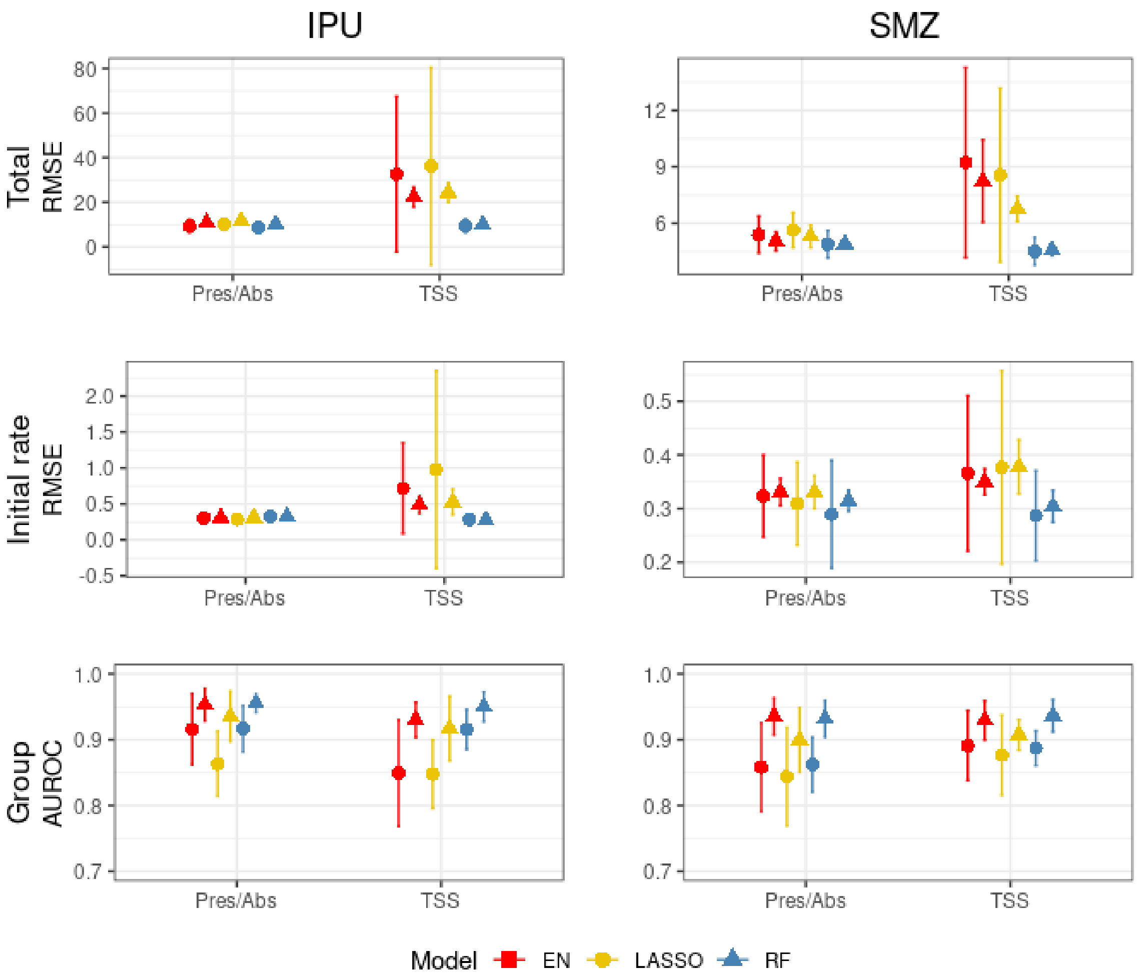


Figure 3: **RMSE and AUROC for all phenotypes and statistical methods**. RMSE (Root Mean Square Error) for Total and Initial rate; AUROC (Area Under the Receiver Operating Characteristic curve) for degrading groups are shown for elastic net regression (EN), LASSO, and random forest (RF) on relative abundance OTUs data (TSS) and presence Absence of same OTUs (Pres/Abs). All predictions were assessed by 10-fold CV with shuffling between 8 runs, and only results on test set are shown. Errors bars stand for standard error of the mean.

### 4 Total Sum Scaling distribution of degrading communities

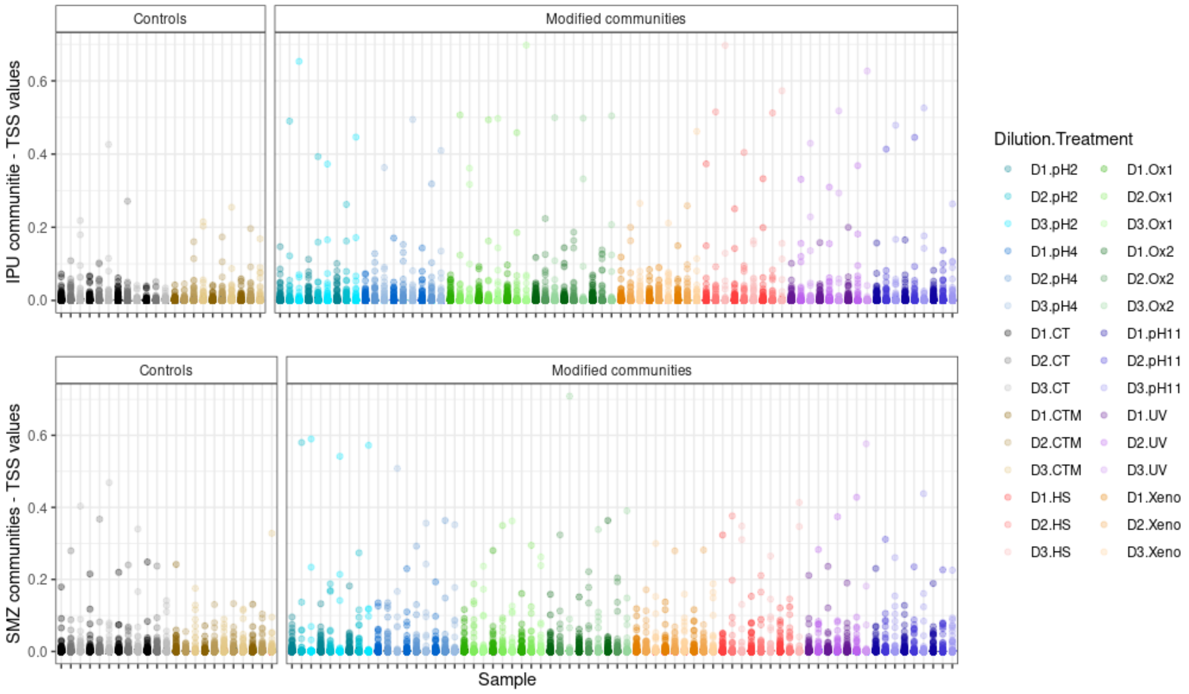


Figure 4: **Total Sum Scaling (TSS) of OTUs from IPU (above) and SMZ (below) degrading communities.** Biocidal treatments are shown by colours and dilutions by colour gradient.

### 5 Examples of predicted classification outcome

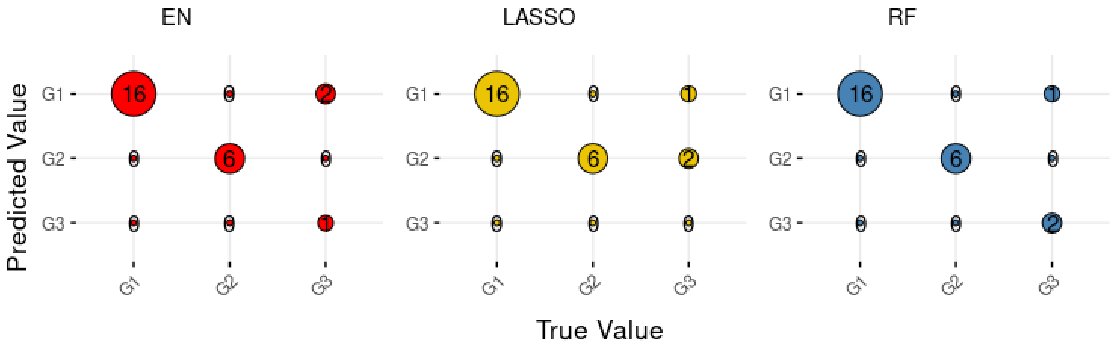


Figure 5: **Example of true to predicted classification for IPU degrading groups.** Three models on PresAbs data, testing set, run 5. Samples are placed depending on their true mineralisation group membership (x axis) and their predicted classification (y axis). Cell size are proportional to the number of samples, which is stated in the cells.

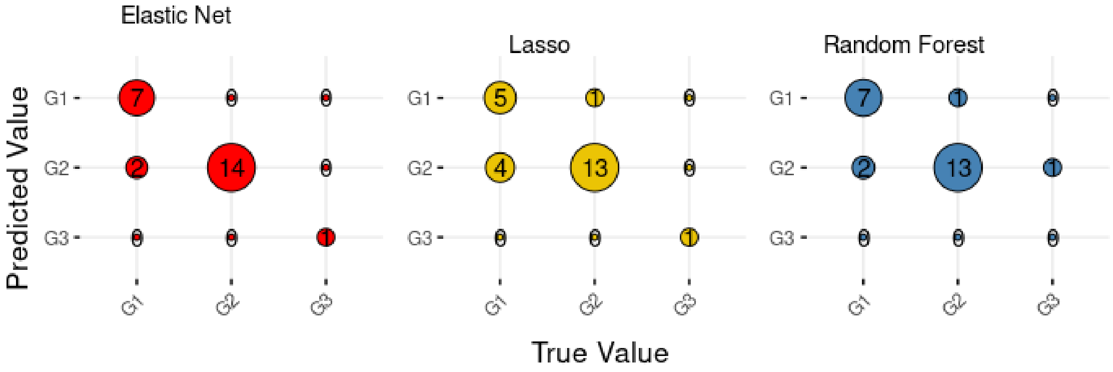


Figure 6: **Example of true to predicted classification for SMZ degrading groups.** Three models on TSS data, run 6. Samples are placed depending on their true mineralisation group membership (x axis) and their predicted classification (y axis). Cell size are proportional to the number of samples, which is stated in the cells.

#### 6 Hyperparameters of prediction methods

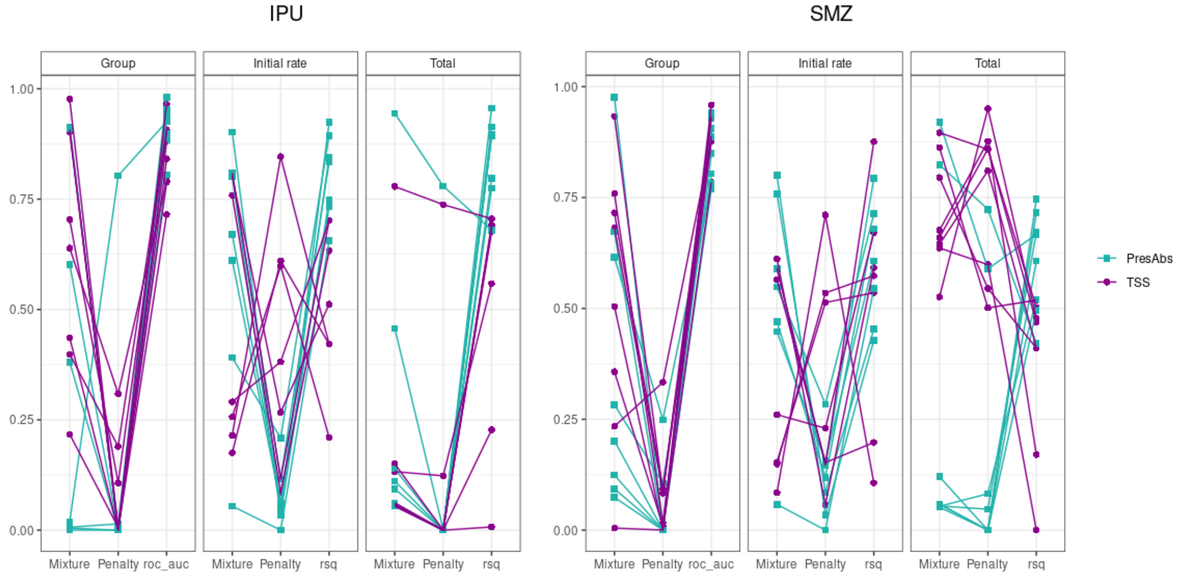


Figure 7: **Hyperparameters values and performances of prediction and classification by Elastic Net for 3 phenotypes and 2 data transformation.** Performances of prediction (Initial rate and Total) are measured by RMSE, performances of classification (Group) by AUROC. The hyperparameters are the mixture : low values for a strong selection of variables (equivalent to LASSO when equal to 0), high value (max 1) for a shrinkage of variables estimates (similar to ridge regression) and the penalty, which controls the degree of regularization (0 is equivalent to simple linear model). There are 8 runs for each phenotype x Data transformation and hyperparamaters are computed by 10-fold CV.

### 7 Variable importance

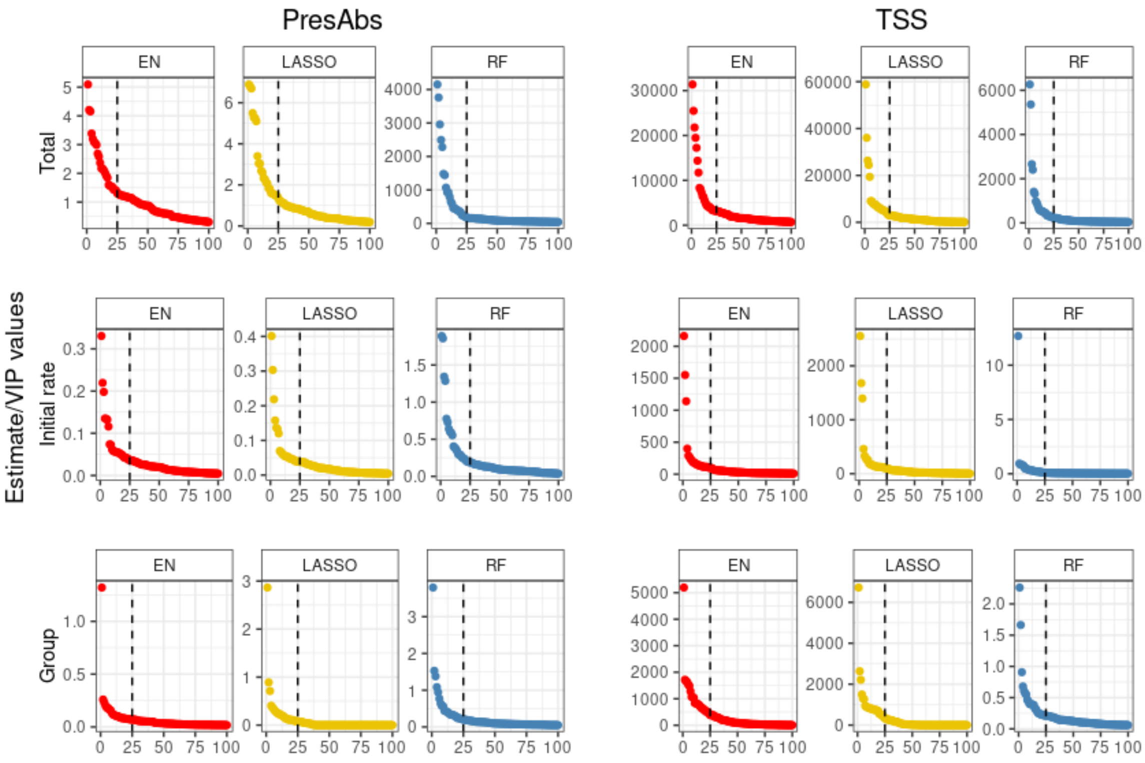


Figure 8: **Variable importance for top 100 OTUs for IPU prediction.** OTUs are shown by decreasing variable importance. Dashed line shows the 25 top OTUs further investigated.

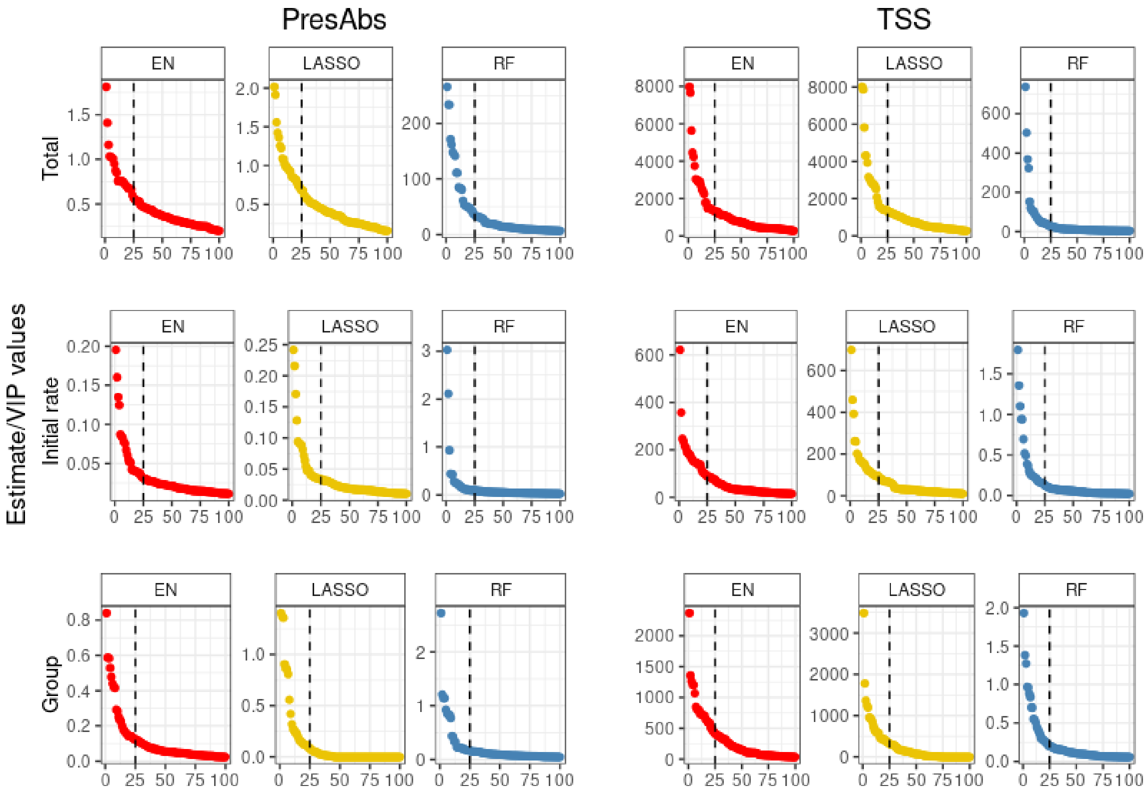


Figure 9: **Variable importance for top 100 OTUs for SMZ prediction.** OTUs are shown by decreasing variable importance. Dashed line shows the 25 top OTUs further investigated.

### 8 Shared important OTUs across data transformation and models

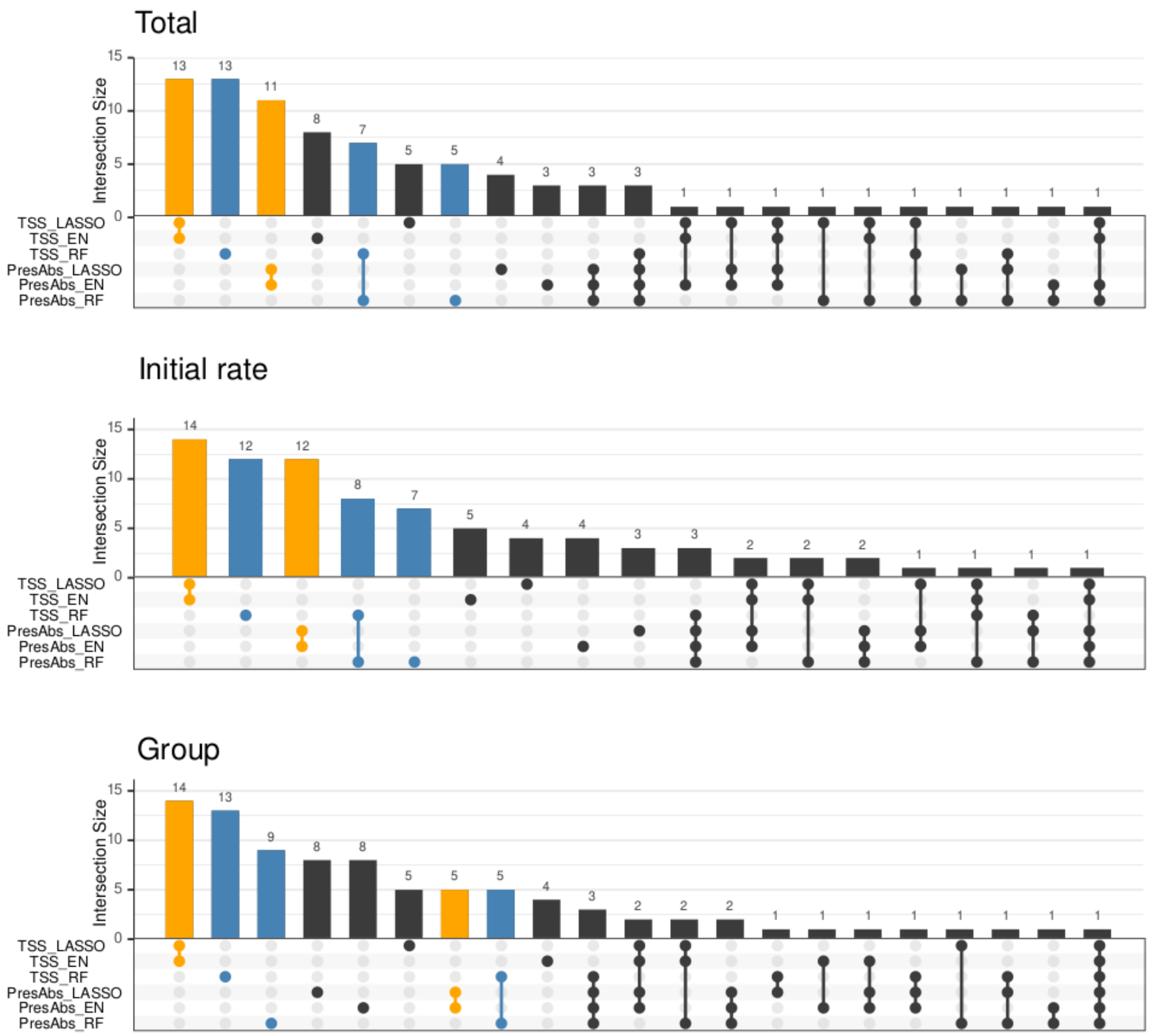


Figure 10: **25 most important OTUs for IPU phenotypes prediction and each data transformation (TSS and PresAbs) coupled with models (LASSO, EN and RF)**. Common OTUs between LASSO and EN are highlighted in gold while OTUs only present in RF are highlighted in blue.

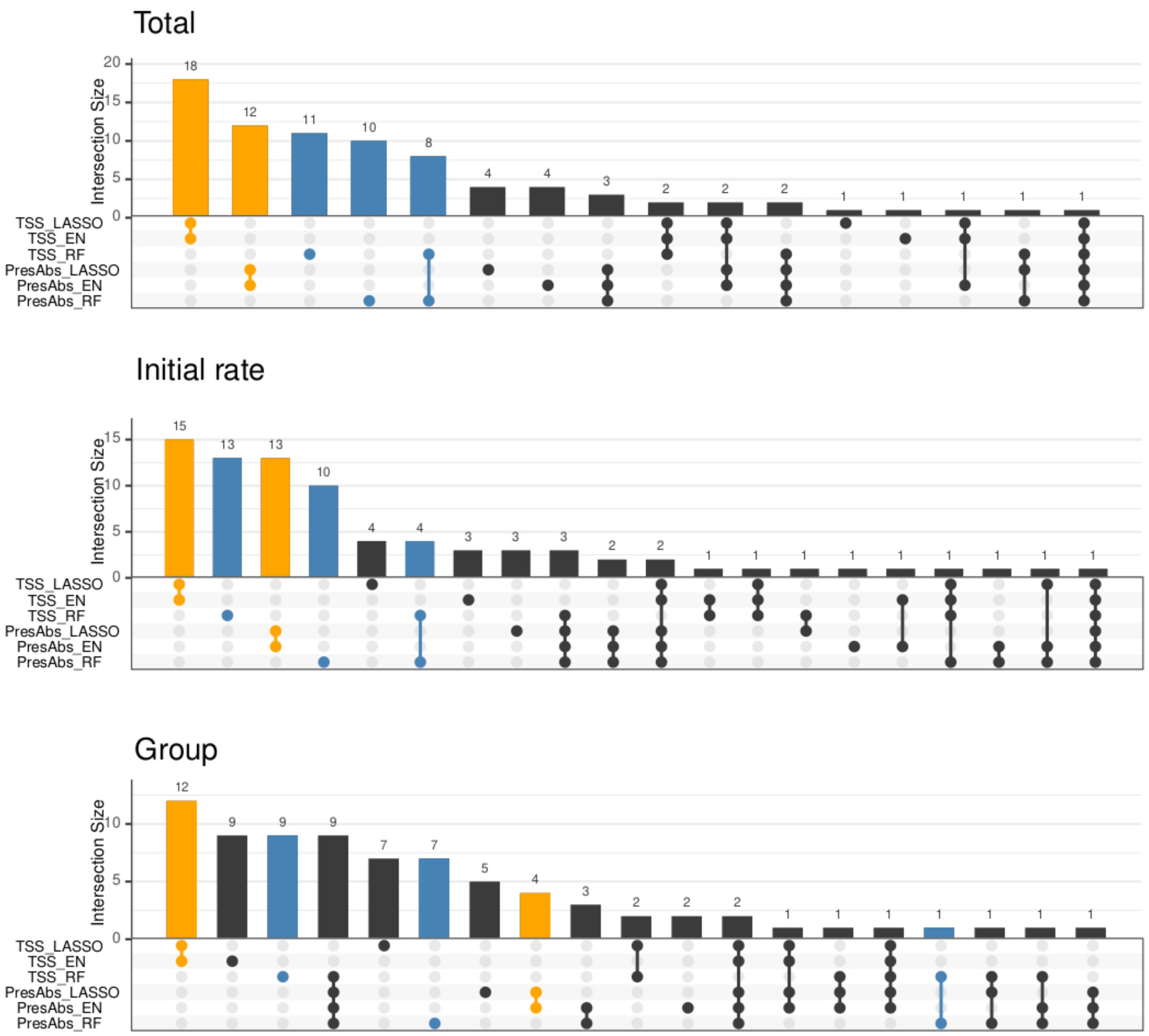


Figure 11: **25 most important OTUs for SMZ phenotypes prediction and each data transformation (TSS and PresAbs) coupled with models (LASSO, EN and RF)**. Common OTUs between LASSO and EN are highlighted in gold while OTUs only present in RF are highlighted in blue.

#### 9 Distribution of OTUs in Louvain groups

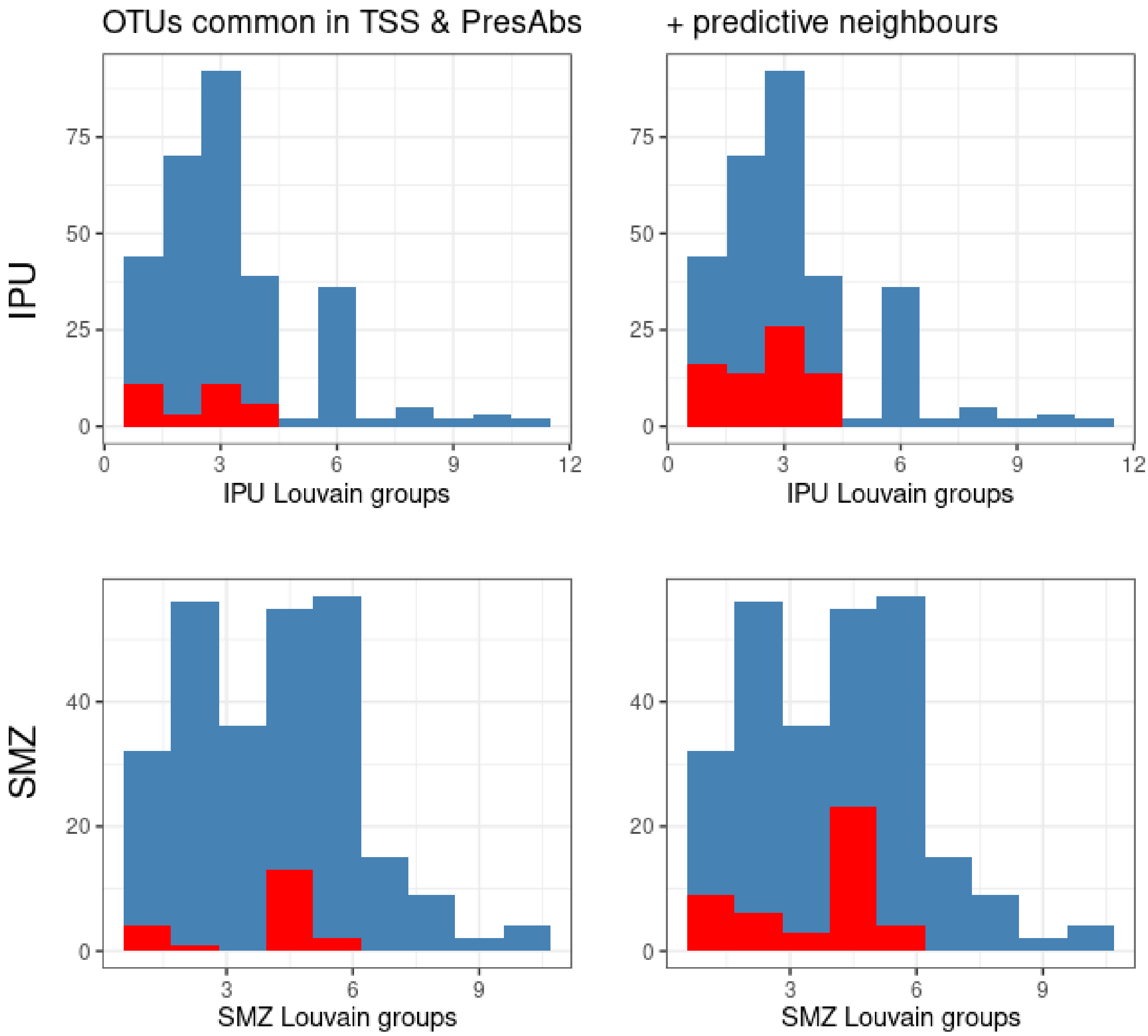


Figure 12: **Distribution of OTUs in Louvain groups & predictive OTUs.** Louvain groups are based on co-occurence network from IPU degrading communities (above) and from SMZ degrading communities (below). OTUs with predictive value both in PresAbs and TSS data transformation are highlighted in red (left). Same ones plus their direct co-occuring neighbours with predictive value in either TSS or PresAbs are represented in red (right)
